# Proteomic insights into the photobiology of the Hawaiian rice coral *Montipora capitata* in response to decreased light intensity

**DOI:** 10.64898/2025.12.02.691798

**Authors:** Tristan Permentier, Hugo Ducret, Emma Timmins-Schiffman, Helena F. Willard, Sophie-Luise Heidig, Christopher R. Suchocki, Robert J. Toonen, Brook L. Nunn, Marc Kochzius, Jean-François Flot

## Abstract

Reef-building corals are sessile marine organisms that inhabit a wide range of light habitats along depth gradients. As coral biology is often studied in the context of global change and changing temperatures, knowledge gaps persist in our understanding of the molecular and cellular pathways involved in the responses to other factors than temperature, such as light intensity, which decreases exponentially in the water column and gradually changes the environment. To fill this gap, we tested the response of the Hawaiian rice coral *Montipora capitata* to decreased light intensity in a field experiment in Kāne’ohe Bay, O’ahu, Hawai’i, using Data-Independent Acquisition (DIA) proteomics. There was a significant effect of light intensity on both the coral and zooxanthellae proteomes. In the *M. capitata* host, 69 proteins differed significantly in abundance between light levels after two years. The 50 proteins identified as significantly more abundant in the control condition were mostly involved in mRNA and RNA processing, pointing toward a positive correlation between metabolic activity, growth rates and increased light levels. The 19 proteins identified as significantly more abundant in the shade treatment were associated with calcium transport and with the structure of key cellular components, such as cell membrane and cytoskeleton. By contrast, zooxanthellae showed only minor changes in protein abundances, with photosynthesis proteins more abundant in the shade treatment and enzymes involved in fatty acid metabolism more abundant in the control treatment. Overall, these findings establish a baseline for our understanding of the cellular and metabolic processes driving *Montipora capitata*’s acclimatization potential to different light intensities.

## Introduction

Coral reefs belong to the most valuable ecosystems on earth, in terms of biodiversity and ecosystem services (Eddy *et al*., 2021; Sing Wong *et al*., 2022). They are currently facing multiple threats, the most critical being the increasingly frequent, thermal-induced coral bleaching events (Bahr *et al*., 2017; Hughes *et al*., 2017; Pei *et al*., 2023). Bleaching is the process in which corals lose their color as they expel the pigmented endosymbionts, revealing the white coral skeleton beneath the symbiont-free tissue (Goreau, 1964). This broad stress response is triggered by various stressors and arises from intricate interactions between biological and environmental factors at the molecular level (Brown, 1997; Suggett & Smith, 2020; Helgoe *et al*., 2024), which highlights the need to elucidate the mechanisms by which the environment affects the coral holobiont (Cunning *et al*., 2015).

Proteins are vital for life’s functions and directly reflect biological activity (Whitford, 2013). The study of an organism’s complete protein set at a specific time, proteomics, can offer insights into the molecular processes at the base of responses to environmental stressors (Nunn *et al*., 2025). Proteins are directly responsible for the biological processes of interest, whereas genes and mRNA go through several regulatory layers before a cellular response can be observed as changes in protein abundance, which makes them more noisy and potentially less accurate (Hegde *et al*., 2003; Xiang *et al*., 2024). As a result, in contrast to genomics or transcriptomics, proteomics offers a more direct view on the current physiological activity of an organism. However, the very high diversity of proteins in a cell means that proteomic analyses usually have a relatively low coverage compared to genomic or transcriptomic datasets, leading to stochastic peptide detection and reduced sampling depth (Cho, 2007).

In the case of scleractinian corals, as the amount of dinoflagellate symbiont proteins is orders of magnitude lower than that of coral host proteins, accurately detecting the proteins of zooxanthellae has been challenging. However, data-independent acquisition (DIA) workflows have improved reproducibility and proteomic depth compared to traditional methods (Collins *et al*., 2017). While proteomics once relied on chromatogram libraries limited to well-studied species, new library-free search techniques now enable peptide detection and quantification even in species lacking reference data, thereby expanding the application of proteomics to non-model organisms, including the coral-symbiont holobiont (Gessulat *et al*., 2019; Demichev *et al*., 2020; Searle *et al*., 2020).

Recent advancements in proteomics have enabled great advances in coral research. For example, Pei *et al*. (2023) investigated the effects of benzo(a)pyrene on *Montipora digitata*, uncovering how this pollutant disrupts coral polyps’ antioxidant capacity, inducing apoptosis in zooxanthellae by suppressing superoxide dismutase 2 (SOD2)-related proteins. Further, Petrou *et al*. (2021) conducted broad proteomic analysis on thermally stressed *Acropora millepora*, revealing crucial differences in host-symbiont responses to elevated temperatures and identifying host glutamine synthetase (GS) and vacuolar-type ATPase (V-ATPase) as two candidate indicators of symbiosis breakdown. Similarly, by using shotgun proteomics on *Montipora capitata*, Axworthy *et al*. (2022) unveiled distinct metabolic pathways in bleached coral tissue and intraskeletal compartments, providing novel insights into the causes and pathways of coral bleaching.

Such proteomic studies are critical to our understanding of how corals respond to environmental cues, yet it has only been addressed in a narrow array of conditions, mostly in a context of rising temperatures (Axworthy *et al*., 2022; Nunn *et al*., 2025; Timmins-Schiffman *et al*., 2025) while paying little attention to changing light intensities. In contrast to the relatively narrow range of temperatures at which they can be found (Wells, 1957), corals can occupy a wide variety of light habitats (Kahng *et al*., 2019; Muir & Pichon, 2019; López-Londoño *et al*., 2022). Between the upper and lower limits of their depth range, light intensity decreases exponentially (Gordon, 1989), which creates distinct light habitats. As such, the physiology of corals inhabiting mesophotic (dark) and shallow (bright) habitats are known to significantly differ (Johnston *et al*., 2022; Rodriguez-Casariego *et al*., 2024). Before elucidating which pathways are triggered under stress, it is vital to understand the variety of physiological conditions corals can exhibit in different, naturally-occurring, non-stressful, light environments. For instance, corals growing in shallow environments such as nearshore lagoons are often metabolically more active (Roth, 2014), but contain significantly more proteins related to DNA repair or antioxidant activity (Falkowski & Chen, 2003; Malik *et al*., 2021), due to high light intensity inducing photodamage and increased protein turnover (López-Londoño *et al*., 2022). Conversely, algal symbionts of corals growing in darker environments often have higher photosynthetic yields and contain more photopigments (Falkowski, 1981; Hochberg *et al*., 2003; Roth, 2014), which optimizes absorption of incident light, and higher heterotrophic activity is often observed in the coral host (Heikoop *et al*., 1998; Maier *et al*., 2010; Padilla-Gamiño *et al*., 2019). Similarly, calcification rates are usually decreasing with lower light levels (Rinkevich & Loya, 1984; Allemand *et al*., 2011; Bahr *et al*., 2017). This phenomenon is called light-enhanced calcification (Chalker &Taylor, 1975), and is a consequence of higher photosynthetic yields at higher light levels. All of these cited mechanisms are directly dependent on light availability, which drives photosynthetic yield (DiPerna *et al*., 2018) and determines how much energy can be allocated to different cell functions.

To date, no study has looked at the impact of light intensity on the proteome of hard corals. These variations in responses can most easily be observed in species that are capable of acclimatizing to a wide range of environmental gradients, such as the depth-generalist Hawaiian rice coral *Montipora capitata* (Veron, 2000). *M. capitata* displays an array of physiological responses that make it an excellent model organism to investigate the acclimatization potential of reef-building corals (Grottoli *et al*., 2006; Drury *et al*., 2022). A striking example of its acclimatization potential is the shift in morphology it presents depending on the light environment, with branching morphologies prevalent at high light and plate-like morphologies prevalent at low light (Veron, 2000; Padilla-Gamiño *et al*., 2013). In this study, we used data independent acquisition (DIA) proteomic analysis to investigate which metabolic functions are most impacted by changing light intensities in *M. capitata* corals and their photosymbionts. We expected to identify differentially abundant proteins between high-light adapted and low-light adapted colonies. In the control treatment, we expected to observe a higher metabolic activity (Roth, 2014), with potential signs of oxidative damage (López-Londoño *et al*., 2022). In the low-light treatment, we expected that the limitation of light may induce a compensatory increase in proteins related to photosynthesis (Roth, 2014), heterotrophic feeding (Heikoop *et al*., 1998) and/or calcification-related processes (Moya *et al*., 2008). All in all, we aimed to establish a baseline at the molecular level for the comparison of shade-adapted and high-light-adapted corals.

## Methods

### Experimental design

The experiment was set up on November 24, 2021, at the HIMB coral nursery, located off the island of Moku o Lo’e within Kāne’ohe Bay, O’ahu, Hawai’i (USA). Six platforms were constructed as follows: three remained under ambient light conditions, and three were shaded using 6’x8’ black mesh screens with 73% shading efficiency (EZ corners, USA). All platforms were built using identical materials and deployed simultaneously to promote consistency in the fouling community. Racks were positioned at a depth of two meters within the floating coral nursery. The nursery is protected against surface currents by its enclosed structure and its location on the leeward side of Moku o Lo’e, ensuring low and uniform water motion around the platforms. HOBO loggers (Onset, USA) were installed on each platform to continuously monitor light intensity and water temperature at 5-minute intervals from the start of the experiment (see Ducret *et al*. (2024), Supplementary Figure S1). Nine *Montipora capitata* parent colonies growing under ambient conditions in the coral nursery were fragmented into six pieces each (n = 54). After fragmentation, the corals were allowed a three-week recovery period under ambient conditions before being moved to the shading treatment or kept under ambient conditions. Each of the experimental platforms received one fragment from each of the parent colonies, which was randomly placed in a 3 x 3 grid arrangement, resulting in triplicates for all nine parental colonies under both light conditions.

### Sample collection

Just before sampling, the maximum quantum yield of each coral colony (F_v_/F_m_ ratio) was measured using a MINI-PAM II (Pulse-Amplitude Modulated, WALZ, Germany) with settings matching the manufacturer’s instructions (MI: 3, Gain: 6, DAMP: 2, SI: 8, SW: 0.8 s). For each colony, we conducted n = 3 measurements of the F_v_/F_m_ ratio to take into account intra-colonial variability.Sample collection took place at noon on February 8th, 2024, two years and two months into the experiment. Although there was some fragment mortality during the experiment, we were able to retain corals from six parental colonies (designated colonies from A to F) for our proteomic analysis (Figure 1A). For colony D as well as for colony E, one coral fragment in the shade condition did not survive. To ensure condition-wise triplicates for both colonies, we sampled from opposite sides of one of the two surviving colonies growing in the shade treatment (Fig. 3). Tips of the colonies’ growing edges measuring approximately 1.0 cm were sampled from each colony (n = 36) and transferred into 1.8 ml microcentrifuge tubes. The samples were stored at -80°C at HIMB and then shipped at -190°C in a dry shipper with absorbed N_2_ to the University of Washington in Seattle, USA.

**Figure 1.**
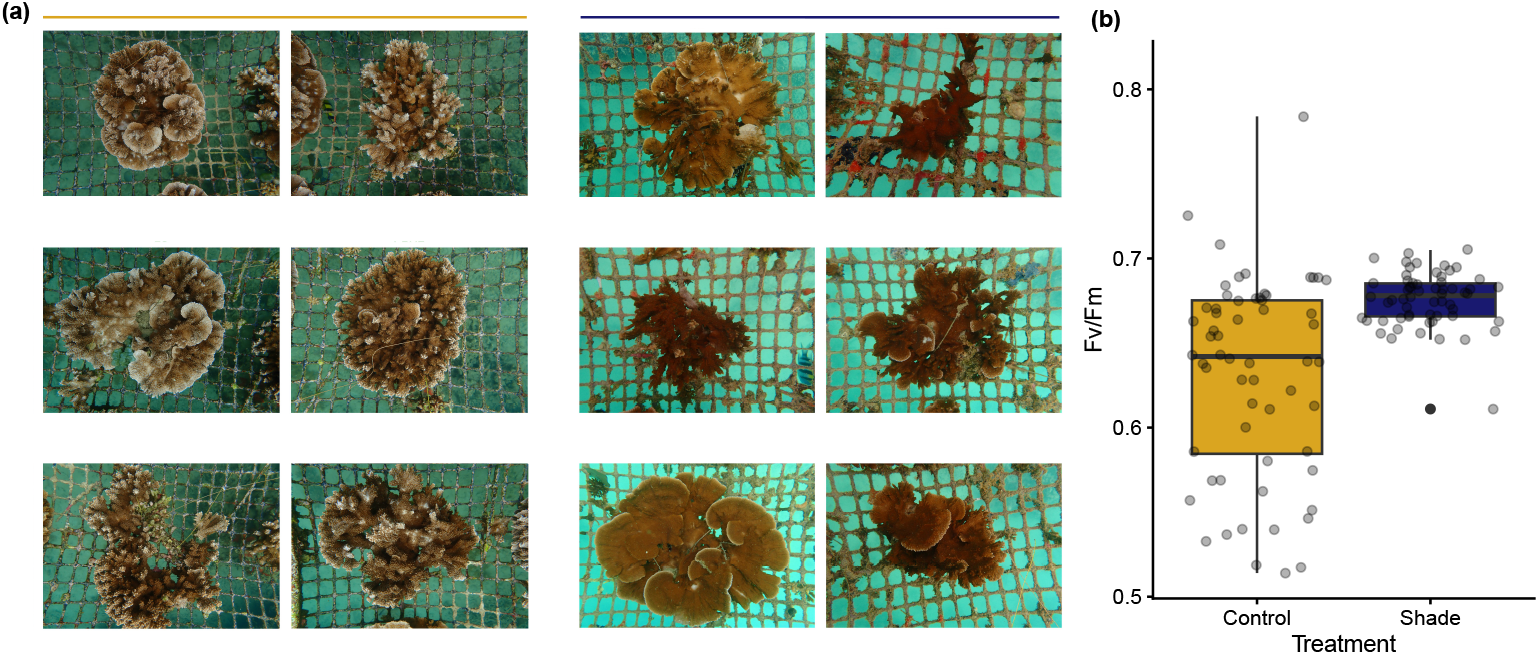
(a) The six *Montipora capitata* coral colonies used for the present study in both treatments. The six pictures below the dark yellow bar (left) correspond to colonies growing at ambient light, while those below the dark blue bar (right) are pictures of colonies grown under 73% shade. The square openings in the mesh in the background are approximately 4x4cm2; (b) Distribution boxplots of the maximum quantum yield of PSII (F_v_/F_m_) values obtained for these corals.

### Protein extraction and digestion

Tryptic peptide samples were prepared using an S-Trap Mini Plate (ProtiFi). Samples (2-3 mm) were homogenized in a Bullet Blender in S-trap sonication buffer (25% SDS (20% stock solution), 5% TEAB, 0.2% MgCl2, 1% HALT protease and phosphatase inhibitors) and adjusted to pH > 7 using 1M Tris HCl if necessary and spiked with 16µl of 100 ng.µl^-1^ enolase and then treated with 0.5 µl of 250 unit.µl^-1^ benzonase nuclease. Yeast enolase was added as an internal control to monitor extraction and digestion efficiency. Samples were then incubated at room temperature for 5 minutes with vortexing. They were then reduced with 1.5 µl of 250 mM tris(2-carboxyethyl)phosphine (TCEP) at 55-60°C for 15 minutes and cooled. Alkylation was performed with 6 µl of 500 mM iodoacetamide (IAA) at room temperature for 30 minutes in the dark. After centrifugation at 13,000 g for 8 minutes, samples were acidified with 8.2 µl of 27.5% phosphoric acid and mixed with 350 µl of S-trap binding buffer (10% TEAB with 90% methanol). The samples were loaded onto the S-trap plate, spun at 1500 g for 2 minutes, washed three times with 200 µl of S-trap binding buffer, and treated with 150 µl methanol/chloroform three times. Trypsin digestion was performed at 47°C for an hour with 4 µg trypsin in 125 µl. Peptides were eluted with 80 µl 50 mM TEAB, 80 µl 0.2% formic acid, and 80 µl 50% acetonitrile/0.2% formic acid, pooled, dried, and resuspended in 100 µl of 2% acetonitrile/0.1% formic acid for LC-MS/MS analysis.

### Liquid chromatography - tandem mass spectrometry (LC-MS/MS)

Prior to conducting mass spectrometry, we used the bradford protocol to calibrate the amount of protein in each sample (He, 2011). Data-independent acquisition (DIA) was carried out using a Lumos mass spectrometer (ThermoFisher). We chose to perform DIA over data-dependent acquisition (DDA) as it yields better coverage of low-abundant proteins compared to DDA (Timmins-Schiffman *et al*., 2025), which was necessary to accurately detect zooxanthellae proteins since they were expected to be less abundant than coral host proteins in our samples. Reverse-phase high-performance liquid chromatography (HPLC) was conducted inline with the mass spectrometer using an EasyLC system (ThermoFisher). Each coral sample was analyzed individually using a wide-window single injection covering the entire 400-1000 *m/z* range, utilizing 8 *m/z* isolation windows with a 90-minute gradient of 3-45% solvent B (80% acetonitrile). For *in silico* spectral library generation, equal quantities of peptides from each sample bundle were pooled, and gas phase fractionation (GPF) was employed to analyze six separate injections across specific *m/z* ranges (400-500, 500-600, 600-700, 700-800, 800-900, 900-1000), each with a narrow 2 *m/z* isolation window. The 130-minute instrument method also included a 90-minute gradient of 3-45% solvent B. Instrument settings for tSIM scans included 120K Orbitrap resolution, an AGC target of 400K, a normalized AGC target of 800%, and centroid data acquisition. For tMSn scans, the settings included a MS2 CID activation time of 10 ms, an orbitrap resolution of 15K, a maximum injection time of 20 ms, an AGC target of 400K and a normalized AGC target of 800%, with centroid data type.

### Protein sequence database construction

We used the protein sequences from the predicted genes in the published *Montipora capitata* genome (Stephens *et al*., 2022). For zooxanthellae proteins, we first identified the zooxanthellae types prevalent in our coral colonies. To do so, vertical branch tip tissue samples of *M. capitata* corals were collected and stored in DNA/RNA shield buffer (Zymo Research, USA) at -20°C immediately after collection until DNA extraction. Amplification of the ITS2 marker gene was performed using the primer pair and PCR cycling conditions suggested by Hume *et al*. (2018). Details about amplification and sequencing of this marker are available in the supplementary material. Based on the results (Supplementary Fig S1), we used the predicted proteome of *Symbiodinium goreaui* (clade C1), now classified under the genus *Cladocopium* (Liew *et al*., 2016; LaJeunesse *et al*., 2018) to build our protein database. We supplemented this database with 125 common laboratory contaminants obtained from the Cambridge Center for Proteomics common Repository of Adventitious Proteins (CCP cRAP) to minimize background noise (https://cambridgecentreforproteomics.github.io/camprotR/articles/crap.html).

### Protein identification and quantification

We utilized DIA-NN 2.0 for protein identification and quantification (Demichev *et al*., 2020). To create a spectral library, we performed an *in silico* tryptic digest of the sequence database using FASTA, allowing for up to two missed cleavages. This process accounted for N-terminal methionine excision and C-terminal carbamidomethylation modifications. The GPF files were utilized as input data for DIA-NN’s deep learning models, which were then employed to predict spectra, retention times, and ion mobilities of the resulting tryptic peptides, thus forming the basis of the spectral library (based on Gessulat *et al*., 2019). The precursor charge was restricted to a range of 1 to 4, with a precursor *m/z* range of 300 to 1800. This spectral library was subsequently used for protein identification and quantification in wide-window coral sample files through heuristic protein inference. The analysis was optimized for high-precision quantification using the QuantUMS strategy, with the match between runs (MBR) feature enabled to enhance identification consistency across different runs. (Kistner *et al*., 2023). DIA-NN automatically performs normalization on protein intensity values to ensure consistent quantification across samples.

### Statistical analyses

To look at the effect of treatment on the F_v_/F_m_ values, we built a linear mixed model with colony as a random effect using the `lmer` function in the `lme4` R package (Bates *et al*., 2015). Differences in relative host/symbiont protein signal intensities between control and shaded colonies were analyzed with a logistic regression computed wit the `glmer()` function of the same package, with colony as random effect. Proteomic variations across all sample types were visualized using non-metric multidimensional scaling (NMDS) plots (Shepard, 1962a,b) via the `metaMDS` function of the `vegan` R package (Oksanen *et al*., 2024). These plots were based on log_2_-transformed protein intensity values, for both algal and coral proteomes. Permutational analyses of variance (PERMANOVA) (Anderson, 2001) were conducted on Bray-Curtis dissimilarity (Bray & Curtis, 1957) matrices to test for the impacts of colony, light levels, or a combination of both on coral and algal proteomes (Borcard *et al*., 2018). Additionally, we looked for potential batch effects associated with S-trap plate variables such as plate row and plate column.

In this study, the differential abundances of host and zooxanthellae proteins were analyzed using linear models (Rao & Toutenburg, 1995) that included light treatment and colony. These linear models were then fitted to the protein expression data using the `lmFit` function of the `limma` R package (Ritchie *et al*., 2015). Models were refined by calculating empirical Bayes statistics using the `eBayes` function. A moderated t-test (Phipson *et al*., 2016) was used to identify differentially abundant proteins in control and shade treatments, with adjustments for multiple testing using the Benjamini-Hochberg method (Benjamini & Hochberg, 1995) to account for false discovery rate (FDR). Proteins were categorized as significantly more abundant in control (log_2_(FC) > 0) or in the shade treatment (log_2_(FC) < 0) based on statistical significance (*p*_adjusted_ *<* 0.05) and a log_2_(FC) threshold of ± 1.5. All statistical analyses were conducted in RStudio v4.5.1 (Ihaka & Gentleman, 1996).

### Biological annotation and functional enrichment analysis

Proteins were annotated using the EggNOG mapper (Cantalapiedra *et al*., 2021) and the UniProt databases (Lussi *et al*., 2023). To search for significantly enriched functions, we used two types of pathway analyses, namely Gene Set Enrichment Analysis (GSEA) (Subramanian *et al*., 2005) and Over-Representation Analysis (ORA) (Ziemann *et al*., 2024). GSEA was employed on the complete protein set to identify functions that were over-represented “as a whole”, while ORA was performed to highlight functions that were over-represented among our differentially abundant proteins only, in order to obtain a more complete picture of the processes occurring in both treatments. The `enricher()` function from the `clusterProfiler` R package (Yu *et al*., 2012) was used to identify significantly over-represented Gene Ontology (GO) terms, while the `topGO` R package (Alexa *et al*., 2006) applied a Fisher’s exact test to extract the top enriched GO terms (Gene Ontology Consortium, 2004). The Gene Ontology hierarchy was visualized and considered in the interpretation of the GO enrichment results using the `showSigOfNodes` function in `topGO`.

## Results

### Fluorometry

We found a significant effect of light treatment on the measured F_v_/F_m_ ratios (Fig. 1B; mixed model, p-value < 0.05), with values of control colonies being, on average, lower than the values of shaded colonies.

### Protein detection

The analysis resulted in the identification and quantification of 8378 proteins in total. Among these, 18 proteins (0.2%) were identified as adventitious (i.e., contaminant) proteins, 6613 proteins (78.9%) were identified from the *Montipora capitata* coral host and 1747 (20.9%) were from the zooxanthellae. Among the 8612 protein groups identified, only 307 (3.56%) were found to contain more than one protein. The relative percentage of protein signals (Fig. 2) was calculated as the total log_2_ protein abundance grouped by organism. The ratio between zooxanthellae and host protein signal intensity did not significantly vary across light conditions (logistic model, p-value > 0.05).

**Figure 2.**
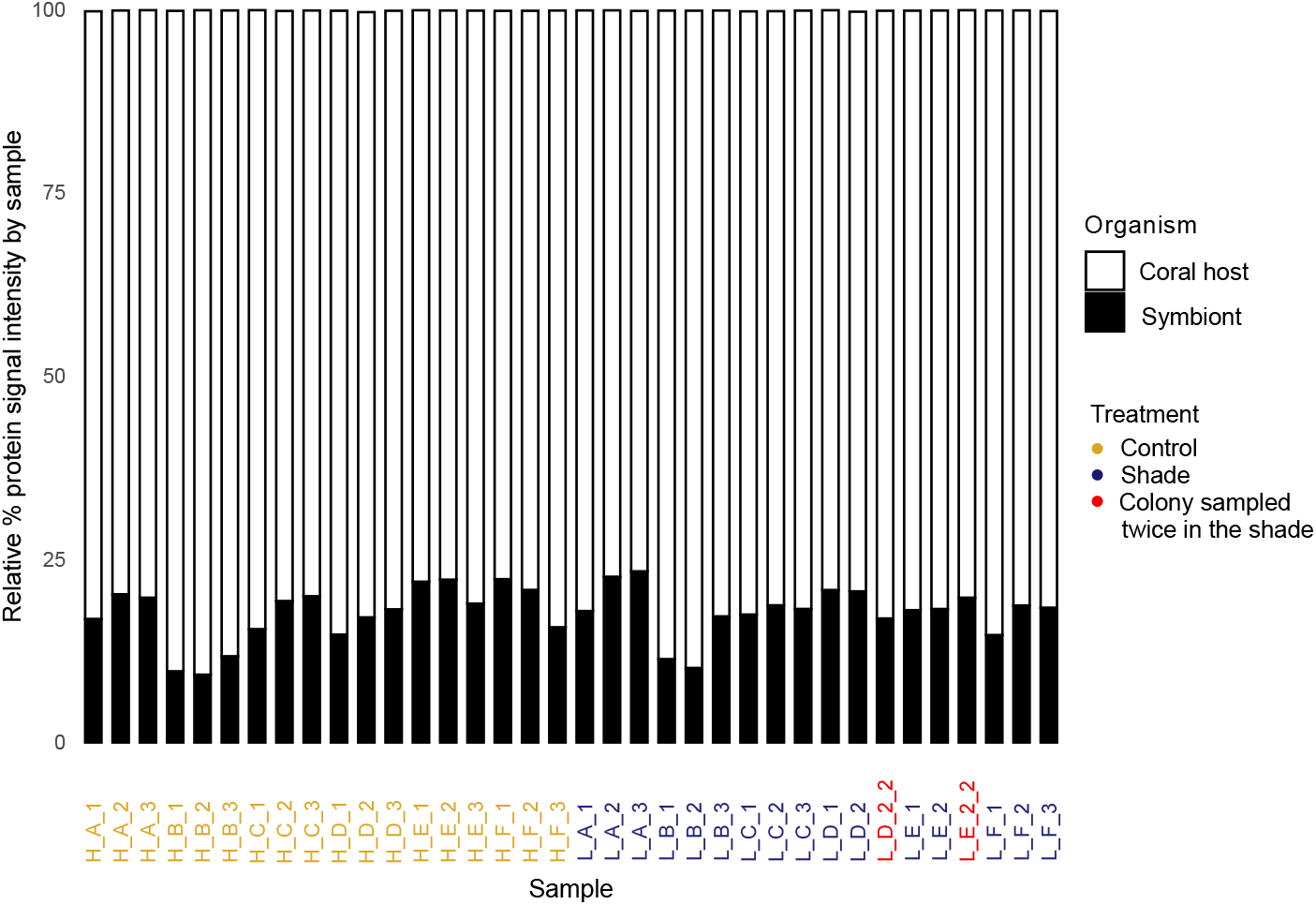
Relative percentage of protein signals detected in samples of *Montipora capitata* attributed to the coral host (white bars) and to the Symbiodiniaceae symbionts (black bars) across samples in the control (yellow labels), shade (dark blue labels), and instances where the same shaded colony was sampled twice (red labels).

**Figure 3.**
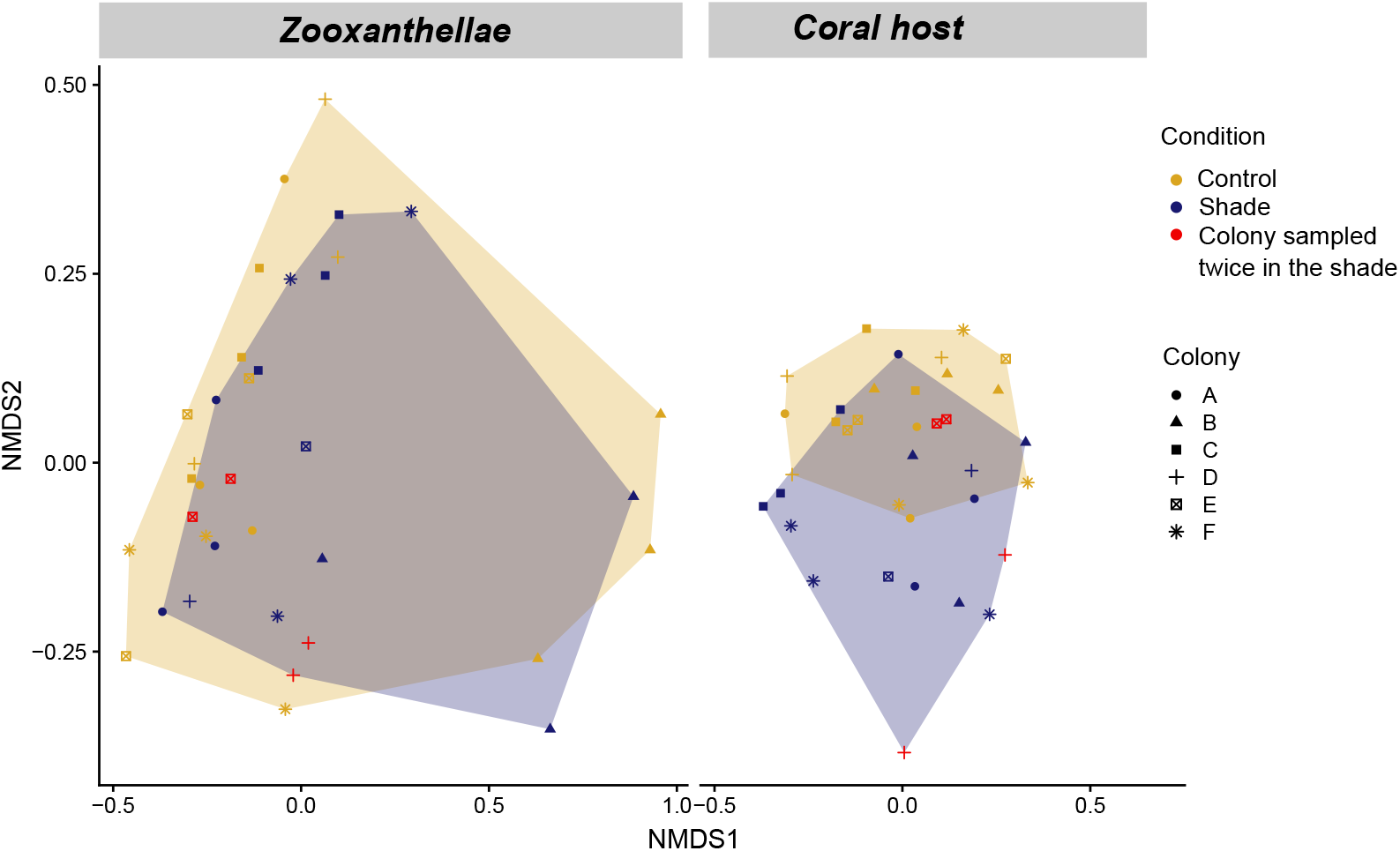
NMDS plots showing variation in coral and zooxanthellae proteomes under different light conditions, with hulls of different colors representing groupings by light level and symbols differentiating colonies. Red markers highlight fragments that were sampled twice.

### Relative protein abundance in hosts and photosymbionts

Within the 6613 proteins identified in coral hosts, 136 (2.0%) were unique to colonies growing in the control treatment, 45 (0.7%) were unique to colonies growing in the shade, and 6595 (97.3%) were found in both treatments (Supplementary Fig. S5). A similar pattern arose with the zooxanthellae proteins, where 95.5% were found in both treatments (Supplementary Fig. S5). We found no batch effect associated with the position of our samples in the mass spectrometer (Supplementary Fig. S2; PERMANOVA, p-value_row_ > 0.05, p-value_column_ > 0.05). For coral hosts, we found a significant effect of light intensity (PERMANOVA, p-value < 0.05), and colony (PERMANOVA, p-value < 0.05) on relative protein abundance. Post-hoc tests highlighted colony C as significantly distinct from four other colonies, and colony A and F as being significantly distinct (Supplementary Fig. S3. For the symbionts, there was no effect of light intensity on relative protein abundance (PERMANOVA, p-value > 0.05). However, we found a colony effect on zooxanthellae proteome composition (PERMANOVA, p-value < 0.05). We identified colony B as an outlier (Supplementary Figure S4). Preliminary ITS2 sequencing results indicated *Durusdinium* sp. dominated the zooxanthellae communities originating from colony B (Supplementary Fig. S1), while the five others were dominated by *Cladocopium* sp. symbionts.

### Differentially abundant proteins and functions

The results of the differential abundance analysis are shown in Fig. 4, revealing 69 significantly differentially abundant proteins in *Montipora capitata*, with 19 elevated in the shade treatment and 50 elevated in the control treatment (moderated t-test, *p*_adjusted_ *<* 0.05 and |log_2_ FC| *>* 1.5). In the significantly differentially abundant proteins of the shade treatment, we identified ATP2B1, a Ca^2+^ transport ATPase, as well as LAMC3 and two MFGE8 orthologs, all of which are extracellular matrix proteins. Among these proteins, our ORA mostly identified significantly enriched GO terms in calcification-related processes (Fig. 6: extracellular space, GO:0005615; P-type calcium transporter activity, GO:0005388; calcium channel regulator activity, GO:0005246).

**Figure 4.**
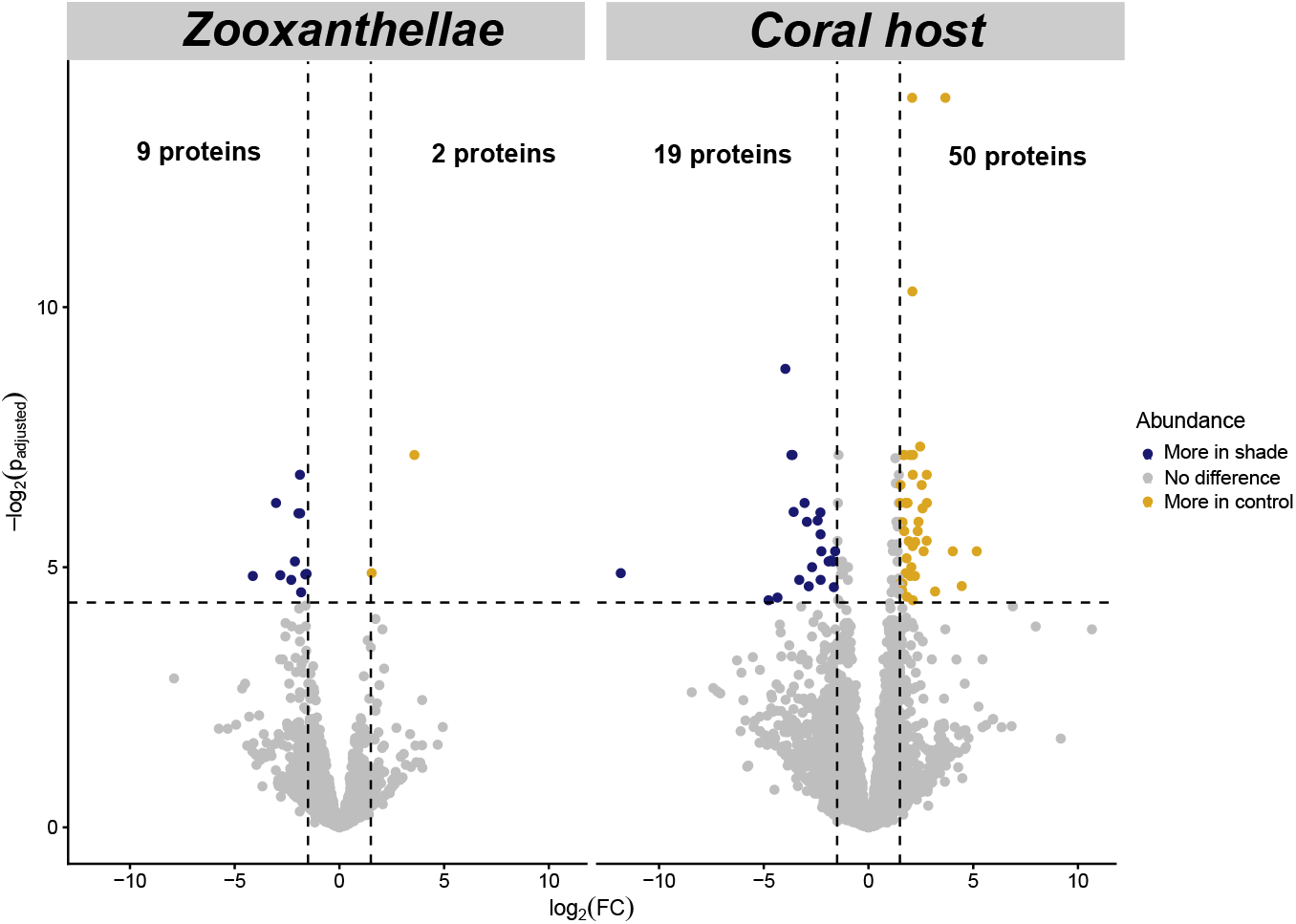
Volcano plots showing the results of differential protein abundance analysis in the *Montipora capitata* coral host and in its Symbiodiniaceae symbionts under control and and shaded conditions. Dashed lines represent significance thresholds (*p*_adjusted_ *<* 0.05 and | log_2_ FC| *>* 1.5).

In contrast, the proteins that were less abundant in the shade treatment encompass several involved in cytoplasmic translation, such as RL18, EIF3A, and RL31 (Fig. 4, Fig. 6). Additionally, we found proteins involved in RNA-related processes, such as alternative mRNA splicing (SRSF6, SRS1B) and mRNA binding (LARP6, THOC4). The SURF1 and QRC9 proteins, related to the inner membrane of the mitochondrion, were also significantly differentially abundant (Fig. 6), as well as the SSB protein involved in DNA repair and replication (Fig. S6). Lastly, PNPO and PROSC are critical for maintaining vitamin B6 homeostasis and were significantly more abundant in the control treatment (Fig. 4, Fig. 6).

Ten significantly differentially abundant Symbiodiniaceae proteins were identified, with nine elevated in the shade treatment and one elevated in the control treatment (Fig. 4, Supplementary Fig. S6; moderated t-test, *p*_adjusted_ *<* 0.05 and |log_2_ FC| *>* 1.5). In the shade treatment, we found that psbU (photosystem II stabilization) and GPR180 (cellular signaling) were significantly more abundant. In addition, the FCP and FCPB proteins were significantly differentially abundant, and are involved in the response to light stimuli. In the control treatment, only ACAC (acetyl-CoA carboxylase, GO:0003989), involved in fatty acid biosynthesis, was detected as a significantly more abundant protein (Fig. 4, Supplementary Fig. S6).

### Gene Set Enrichment Analysis (GSEA)

In the total set of proteins, we identified 293 proteins involved in the structure of the extracellular region (i.e. the cytoskeleton), as well as 246 involved in the extracellular matrix in the shade treatment (Fig. 5). We also identified 192 proteins involved in calcium ion binding, as well as 38 in the binding of integrins, which are transmembrane receptors.

**Figure 5.**
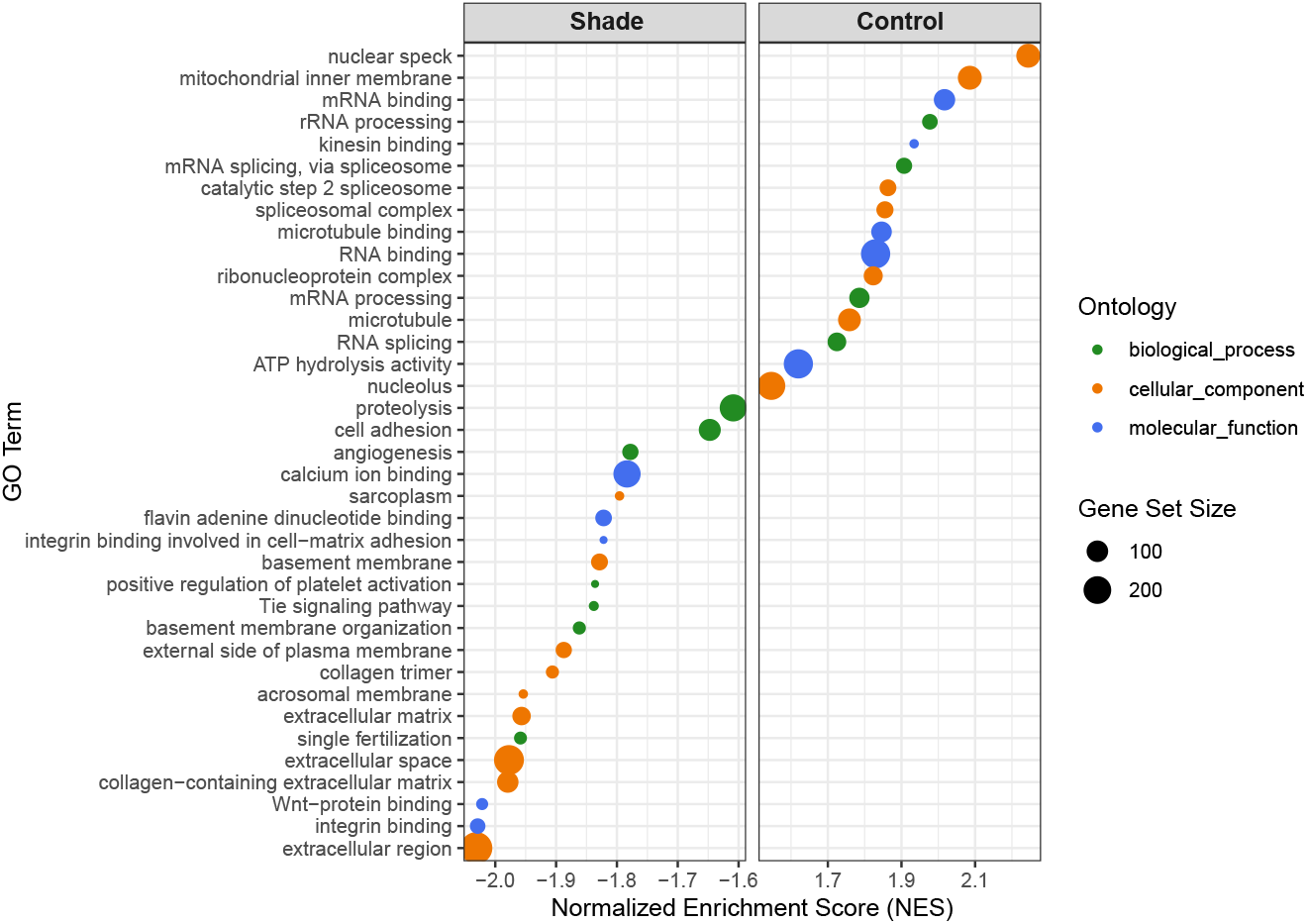
Gene Set Enrichment Analyis (GSEA) of *Montipora capitata* proteins in the control and shade treatments. This analysis was performed on the entire protein dataset. All the Gene Ontology (GO) terms displayed here had an adjusted p-value < 0.05.

**Figure 6.**
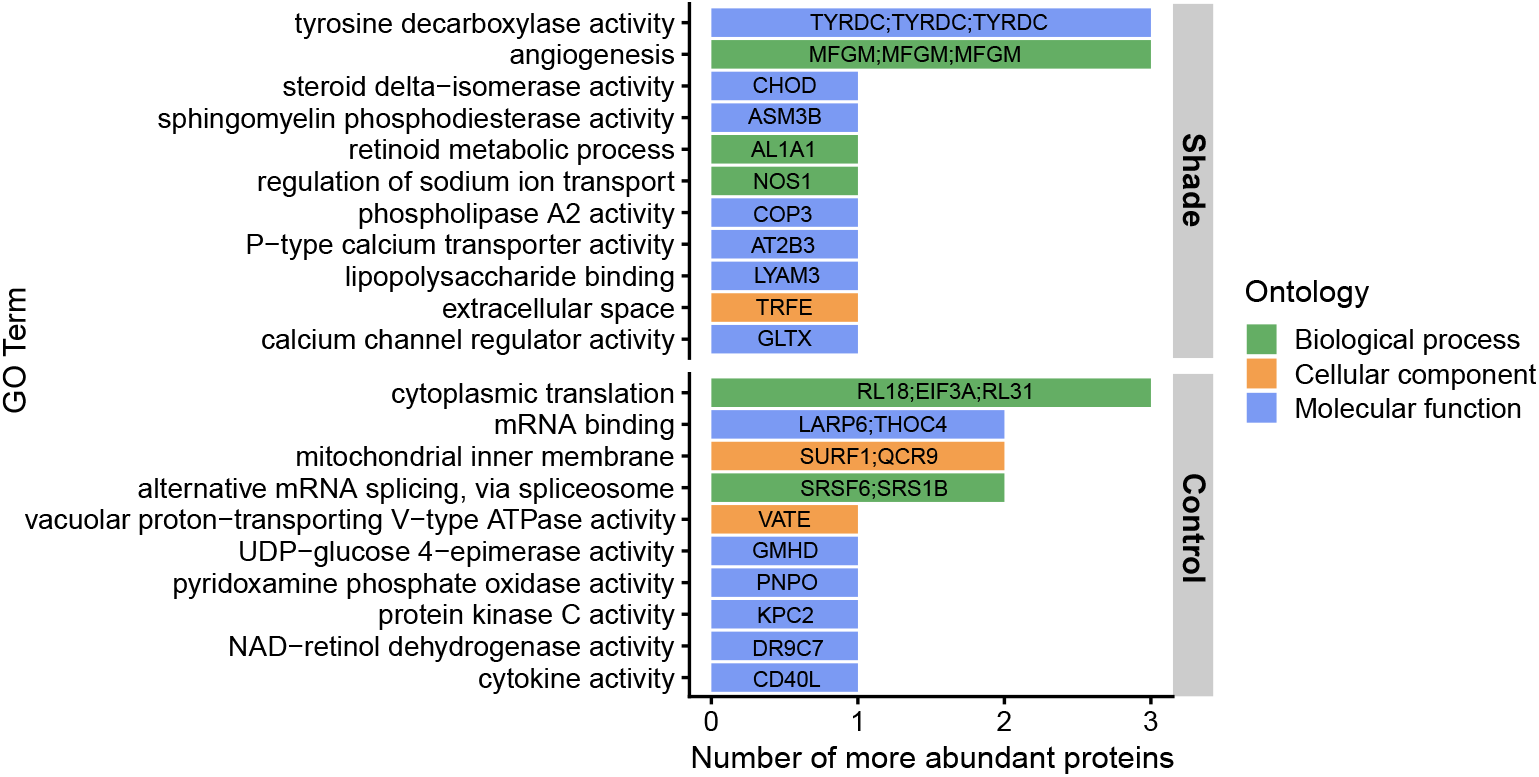
Over-Representation Analysis (ORA) showing the significantly enriched Gene Ontology (GO) terms of control and shaded *Montipora capitata* corals in our study (Fisher’s exact test, p-values < 0.05). This analysis was performed on the subset of differentially abundant proteins identified in Fig. 4. The ontology of the different GO terms (Biological processes, Cellular component, Molecular function) was retrieved from the Uniprot database, and corresponds to different types of functions.

In the control treatment, our GSEA revealed 134 proteins involved in the nuclear speck, and 134 others involved in the mitochondrial inner membrane. In addition, 232 proteins were involved in RNA binding, 98 in mRNA binding, and 74 in microtubule binding (Fig. 5). Lastly, we found 231 proteins involved in the hydrolysis activity of the ATP. The small numbers of differentially abundant zooxanthellae proteins detected did not allow for GSEA or ORA, as the statistical power was too low.

## Discussion

### Evidence of proteomic plasticity in coral hosts

In this study, we used proteomics to investigate how decreased light intensity influences the physiology of both *Montipora capitata* corals and their symbiotic dinoflagellate algae. Our results demonstrate that decreased light intensity has a significant effect on coral host proteomes, with distinct physiological responses observed between colonies growing under 73% shade and ambient light (Fig. 3), which provides evidence of “proteomic plasticity” (Timmins-Schiffman *et al*., 2025), a response similar to what has been described by Rodriguez-Casariego *et al*. (2024) on different biological systems. Decreased light intensity resulted in a higher abundance of proteins associated with key structural cell components in the coral host, such as the cytoskeleton, the cell membrane, and the extracellular matrix, as well as with calcium transporters (Fig 4, Fig.6, Fig.5). Conversely, we observed fewer abundance of proteins involved in mRNA binding, protein turn-over and rRNA and mRNA processing (Fig. 6, Fig. 5). However, Symbiodiniaceae showed limited variation in protein expression. For instance, ACAC, involved in acetyl-CoA carboxylation (Huerlimann *et al*., 2015; Haq *et al*., 2017), was the only zooxanthellae protein significantly more abundant in the control treatment (Fig. 4), which highlights its central role in cellular homeostasis. This raises questions about the potential loss of insight from not isolating symbionts prior to mass spectrometry. Although laborious, this step could have enhanced zooxanthellae proteome coverage.

Further, it is important to note that only a few differentially abundant proteins were identified, which contrasts with other coral proteomics studies that often find dozens (Axworthy *et al*., 2022; Nunn *et al*., 2025; Timmins-Schiffman *et al*., 2025). One possible explanation lies in the magnitude of the changes between the tested conditions. In our experiment, both conditions were viable, while most previous studies looked at proteomic responses to severe heat stress. Alternatively, another possible explanation lies in the differing time scales of the experiments. *M. capitata* is known to be a highly plastic species (Grottoli *et al*., 2006; Drury *et al*., 2022), so the 2-year time frame may have provided enough time for these corals to be well acclimatized. As a result, they regained homeostasis, and the few differentially abundant proteins that were identified may be key functional proteins that allow *M. capitata* to persist in either light treatment. By contrast, Ducret *et al*. (2025) observed marked differences in morphology induced by light-driven phenotypic plasticity on the same corals. Colonies in the control treatment exhibited corymbose-like morphologies, while shaded colonies exhibited plate-like morphologies. These observations suggest that, at this time-point, light acclimation is occurring more at the morphological level than at the proteomic level. As proteomes change more rapidly than morphologies, further studies could aim at comparing the temporal proteomic patterns of control and shaded *M. capitata* corals. This may identify more differentially abundant proteins, and shed light on the processes occurring at the onset of light acclimation.

### Down-regulated functions under decreased light intensity

For our shaded corals, our analyses revealed lower levels of proteins associated with RNA binding and processing (Fig. 5). We also detected significantly less abundant proteins in-11 volved in cytoplasmic translation and alternative mRNA splicing (Fig. 4, Fig. 5). As the RNA/DNA ratio has been used as a biochemical growth rate indicator (Meesters *et al*., 2002), this points toward lower metabolic activity and lower growth rates (Meesters *et al*., 2002; Szmant *et al*., 2004). Light is a key regulating factor, shaping the productivity, physiology, and ecology of the coral holobiont (Roth, 2014) and low irradiance is known to slow down growth rates in corals. This is consistent with the analysis performed by Ducret *et al*. (2025) on these same *M. capitata* colonies, which found significantly lower growth rates in the shade treatment. In addition, we found 2.5 times more significantly abundant proteins in the control treatment when compared to the shade treatment (Fig. 4). This also aligns with the findings of Meesters *et al*. (2002), who found higher protein synthesis per cell in colonies under higher light intensities.

High irradiance levels are also known to increase protein degradation, notably, the degradation of the protein D1 in the PSII of the photosymbionts (Hill *et al*., 2011; López-Londoño *et al*., 2022; Helgoe *et al*., 2024). Ultimately, this degradation causes the formation of reactive oxygen species (ROS) by-products, particularly H_2_O_2_, which can move to the host’s cytoplasm and create oxidative damage there (Oakley & Davy, 2018). Here, we found significantly more abundant ROS scavengers such as GSTM3, and the PNPO and PLPBP enzymes in the control treatment compared to the shade treatment. These have been described as involved in antioxidant processes through the regulation of vitamin B6, critical for the production of the antioxidant glutathione (Krueger *et al*., 2014; Dalto & Matte, 2017; Wang *et al*., 2021). The increased abundance of these proteins, combined with the lower F_v_/F_m_ values in the control treatment (Fig. 1b), might suggest that oxidative stress is occurring in control corals due to light-induced protein degradation (Jones & Hoegh-Guldberg, 2001; Helgoe *et al*., 2024), but that this oxidative stress is not present in the shade treatment. In addition, high protein degradation also requires more recurrent protein replacement (López-Londoño *et al*., 2022), so the lower abundance of RNA-related proteins in the shade treatment may also be the result of the lower metabolic cost of protein repair due to photodamage. This corroborates the findings of Malik *et al*. (2021) who found upregulated genes involved in, among others, oxidative phosphorylation, oxidative stress response, and DNA repair in *Stylophora pistillata* corals transplanted from mesophotic to shallow depths, as well as of Rodriguez-Casariego *et al*. (2024) who found upregulated genes involved in replication, recombination and repair in *Acropora cervicornis* corals from shallow reefs compared to deeper reefs.

### Up-regulated functions under decreased light intensity

Our data points toward a significant upregulation of Ca^2+^ transport and V-type proton-transporting ATPase in the shade treatment (Fig. 6, Fig. 5). Furthermore, the two significantly more abundant proteins annotated as MFGE8 orthologs are believed to facilitate the generation of vesicles containing amorphous calcium carbonate at sites of mineralization (Stapane *et al*., 2020; Sun *et al*., 2020). These findings suggest an upregulation of calcification processes in the shade treatment (Allison *et al*., 2011; Barott *et al*., 2015), which is consistent with previous findings on other hard coral species (Malik *et al*., 2021; Gomez-Campo *et al*., 2024), and may be the result of a compensatory mechanism to cope with the reduction of light-enhanced calcification happening in the shade treatment (Moya *et al*., 2008; Allemand *et al*., 2011; Davy *et al*., 2012). Alternatively, these proteins may counterbalance the acidification of the subcalicoblastic and coelenteron mediums associated with decreased light intensity, particularly with a potential accumulation of protons in the sub-calicoblastic space that are not titrated within the coelenteron due to lower photosynthetic rates (Fig. 1B, Moya *et al*., 2008; Davy *et al*., 2012; Ricci *et al*., 2019; Malik *et al*., 2021). Here, the significantly more abundant calcium pump ATP2B1 protein may play a key role in maintaining i) calcium transport, and ii) a high pH in the extracellular calcifying medium (Al-Horani *et al*., 2003; Sandeman, 2008). Similarly, the identified proteins related to the structure of cellular membranes and the regulation of Ca^2+^ from the extracellular calcifying medium may contribute to the facilitation of lipid peroxidation, fostering Ca^2+^ entry in the cells of the calicoblastic layer by making the plasma membrane more leaky (Moya *et al*., 2008; Sandeman, 2008).

In the photosymbionts of corals exposed to the shade treatment, significant up-regulation of psbU and photo-protection protein PCP have been observed in previous studies (Jiang *et al*., 2012; Cooney *et al*., 2024), along with fucoxanthin chlorophyll proteins (FCPb) involved in the response to light stimuli. These FCPb have been shown to be light-harvesting complexes in diatoms Röding *et al*. (2018). The up-regulation of such photosynthetic proteins points toward a compensatory increase in photosystem machinery (Dubinsky *et al*., 1984; Porter *et al*., 1997). This increase corroborates the higher F_v_/F_m_ values obtained in shaded colonies (Fig. 1B), and previous data from these same colonies showing that chlorophyll *a* content is significantly higher in the shade treatment (Ducret *et al*., 2025).

### Inter- and intra-individual variations in proteome composition

Although we found a significant effect of light intensity on the coral host proteomes, colony was found to have the strongest effect on protein abundance (Fig. 3). This is not surprising, given that this species is usually characterized by high inter-individual differences in morphology (Bhagooli, 2003), physiology (Padilla-Gamiño *et al*., 2019), and proteomic responses (Chille *et al*., 2024; Nunn *et al*., 2025; Timmins-Schiffman *et al*., 2025). We also observed substantial differences between samples sourced from the same coral colony, a pattern that emerged in both host and symbiont proteomes Fig. 3). This is consistent with previous findings by Drake *et al*. (2021), who observed variations in transcriptomes among individual polyps within the same colony of *Stylophora pistillata*. These can also be driven by local differences in light environments across the colony surface (Joyce & Phinn, 2002; Ow & Todd, 2010; Wangpraseurt *et al*., 2012; Roth, 2014). Light intensity is known to be very heterogeneous at the surface of coral colonies, with some parts being significantly more exposed to light than others (Wangpraseurt *et al*., 2012). As such, it may be beneficial for future proteomic studies to include intracolonial replicates to draw conclusions that would be more representative of the whole coral colony.

## Concluding remarks

In this study, we showed that decreased light intensity had a significant impact on the physiology of *Montipora capitata* corals. Fragments growing at reduced light levels are characterized by a slower metabolism and lower oxidative damage, and by higher abundances of proteins related to calcification and photosynthesis, to cope with a reduction of light. However, probably because these corals have been acclimated to shade for two years, there was few differentially abundant proteins overall. This suggests that corals regained homeostasis, and that future studies should aim at comparing the short-term effects of decreased light intensity on these corals in order to obtain the complete picture of the acclimation processes.

## Data and code availability

The data and R codes used in this paper are accessible at https://github.com/montiporacapitata/proteomicinsights/tree/main.

## Supplementary material

### Amplification and sequencing of ITS2 marker gene

In the present study, Internal Transcribed Spacer 2 (ITS2) region of the rDNA was amplified using a two-steps PCR (Collard *et al*., 2025) with the pair developed by Hume *et al*. (2018), upon which we added either a forward tail sequence, or a reverse tail sequence, both identical to the products in the ONT barcoding kits. The outer primer set consisted in the same tail sequences for binding, and a unique barcode sequence, retrieved from the ONT native barcoding kits. As 48 different barcode pairs were used, this allowed for pooling up to 48 samples in one Oxford Nanopore Technolgies (ONT) run. The 25µL PCR mix consisted in 12.5µL of PhireTaq Plant Master Mix 2X (ThermoScientific), 1µL of inner primers at 10µM, 0.5µL of DNA template, and volumes were adjusted with Nuclease-Free water (Collard *et al*., 2025). PCR conditions for the first step were as follow: Initial denaturation at 98°C for 4mins, then 15 cycles of denaturation at 98°C for 8sec, annealing at 56°C for 8sec, and extension at 72°C for 10sec. PCR strips were then taken out of the thermocycler, and a unique barcode pair (Forward-Reverse) was added to each well. PCRs were homogenized, re-incubated in the thermocyclers, and the second step of PCR consisted in 25 cycles of denaturation at 98°C for 8sec, annealing at 56°C for 8sec, and extension at 72°C for 10sec (Collard *et al*., 2025). PCR products were checked on agarose gel and pooled in equimolar quantities to constitute the input of the sequencing library. Sequencing libraries were prepared for R10.4.1 flow cells ran on the Mk1C MinION device, using the Ligation sequencing kit SQK-LSK 114 from ONT. The SpotON flow cell was loaded according to the manufacturer’s instructions, and each sequencing run was performed for 24 hours (Carradec *et al*., 2020).

#### Nanopore read analysis

Basecalling was performed through the `guppy basecaller` function of `guppy v6.3.8`. Reads were demultiplexed with the `guppy barcoder` function of `guppy`, and sorted per barcode. We then used the custom pipeline developed by Carradec *et al*. (2020) to analyze the raw ONT data. Reads were mapped to an ITS2 reference database containing 432 sequences of Symbiodiniaceae (Arif *et al*., 2014) using `minimap2` with “map-ont” as options. The reference sequences covered with a minimum of 1% of all sequences were kept for a second round of read-mapping. This second round was performed in order to aggregate reads potentially mis-assigned during the first round of read-mapping (Carradec *et al*., 2020).

**Fig. S1.**
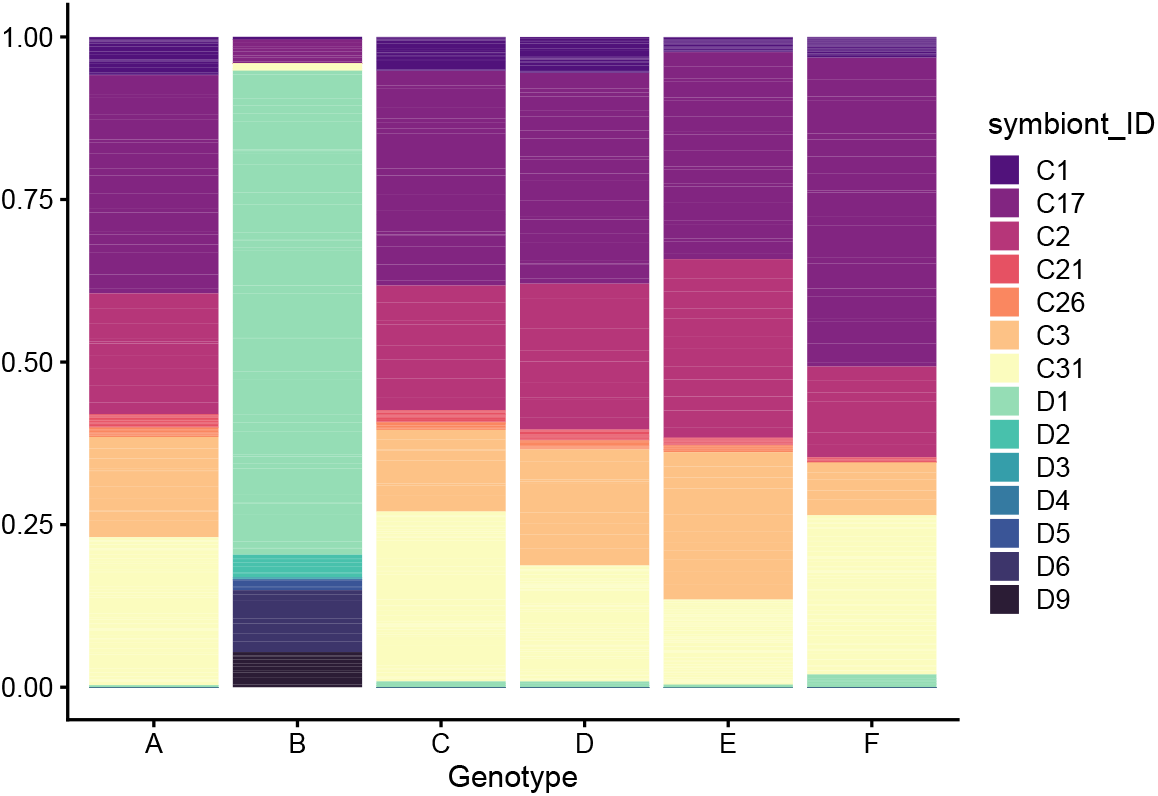
Stacked column plots of zooxanthellae type prevalence with means of relative abundances from samples grouped by colony.

**Fig. S2.**
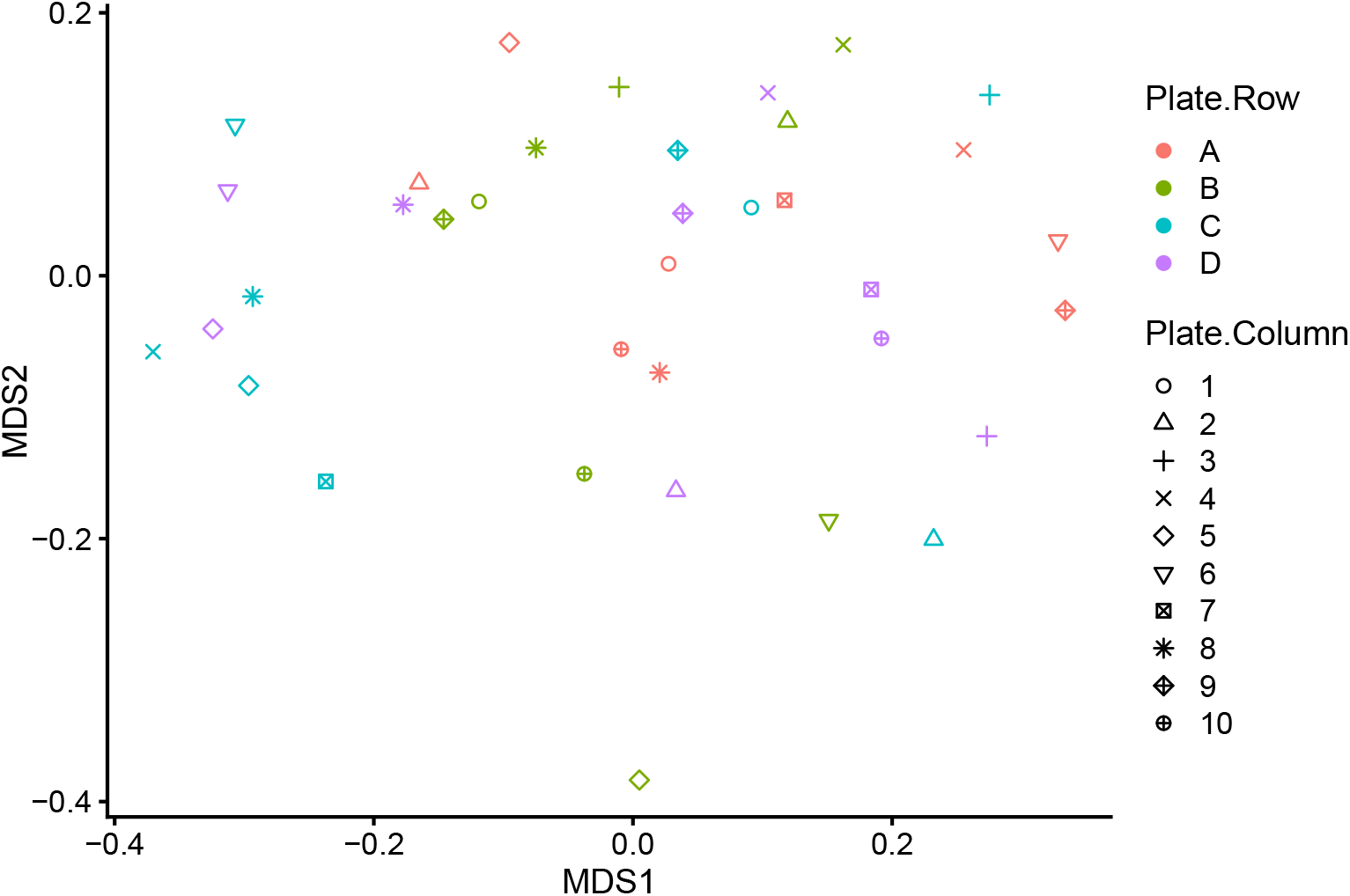
NMDS plot of samples grouped by column and row in the mass spectrometer.

**Fig. S3.**
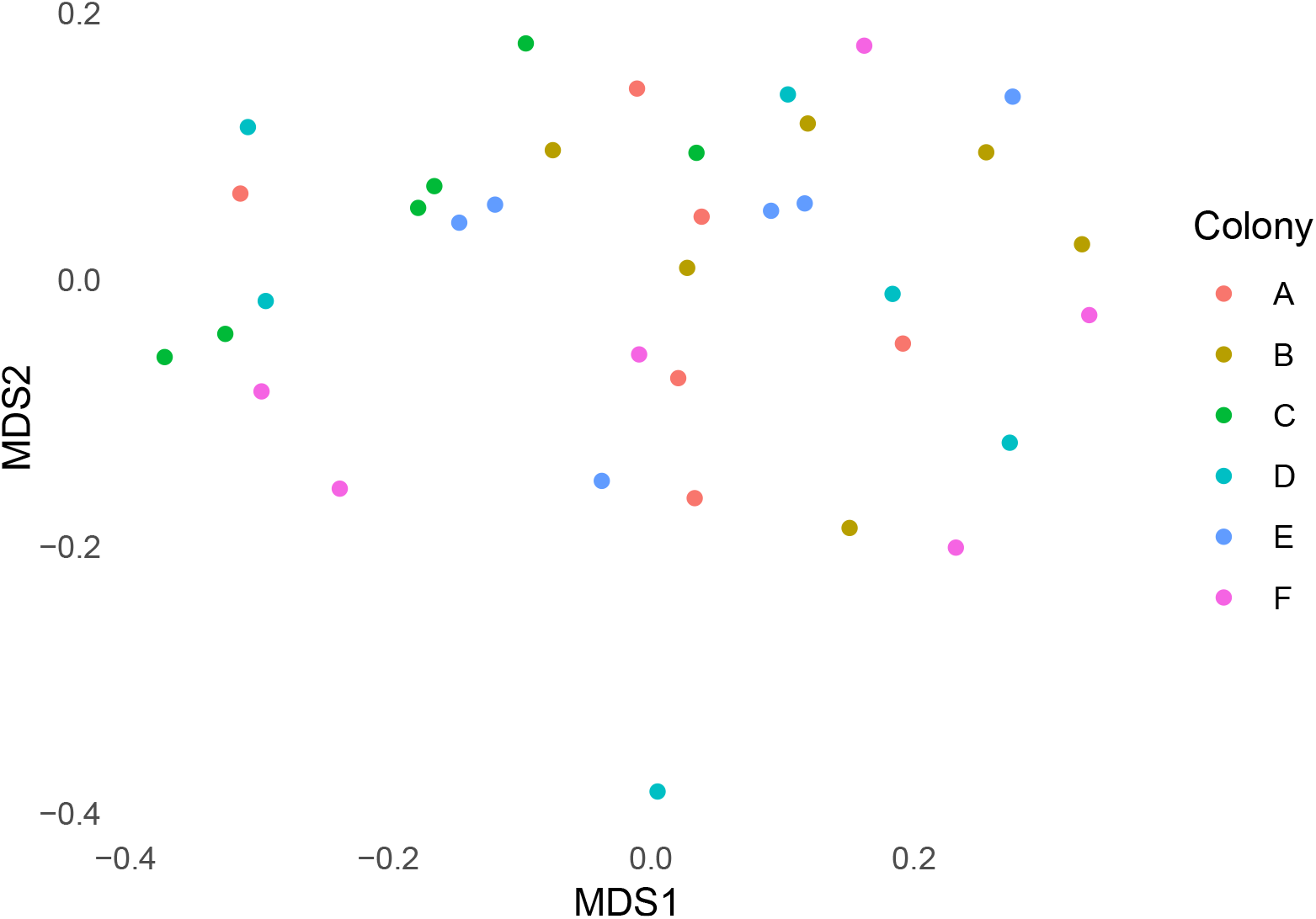
NMDS plot of *Montipora capitata* proteomes grouped by colony.

**Fig. S4.**
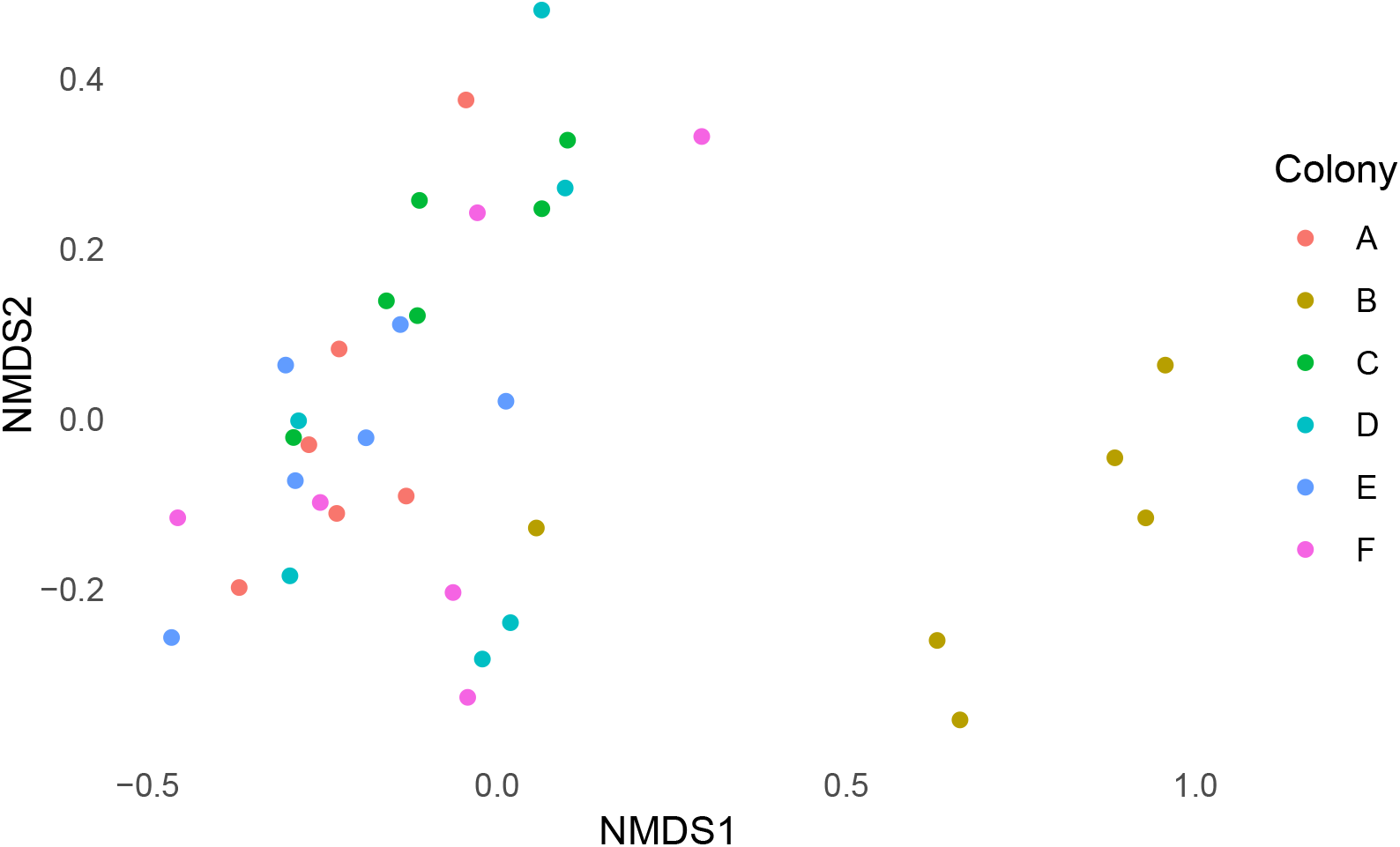
NMDS plot of zooxanthellae proteomes grouped by colony.

**Fig. S5.**
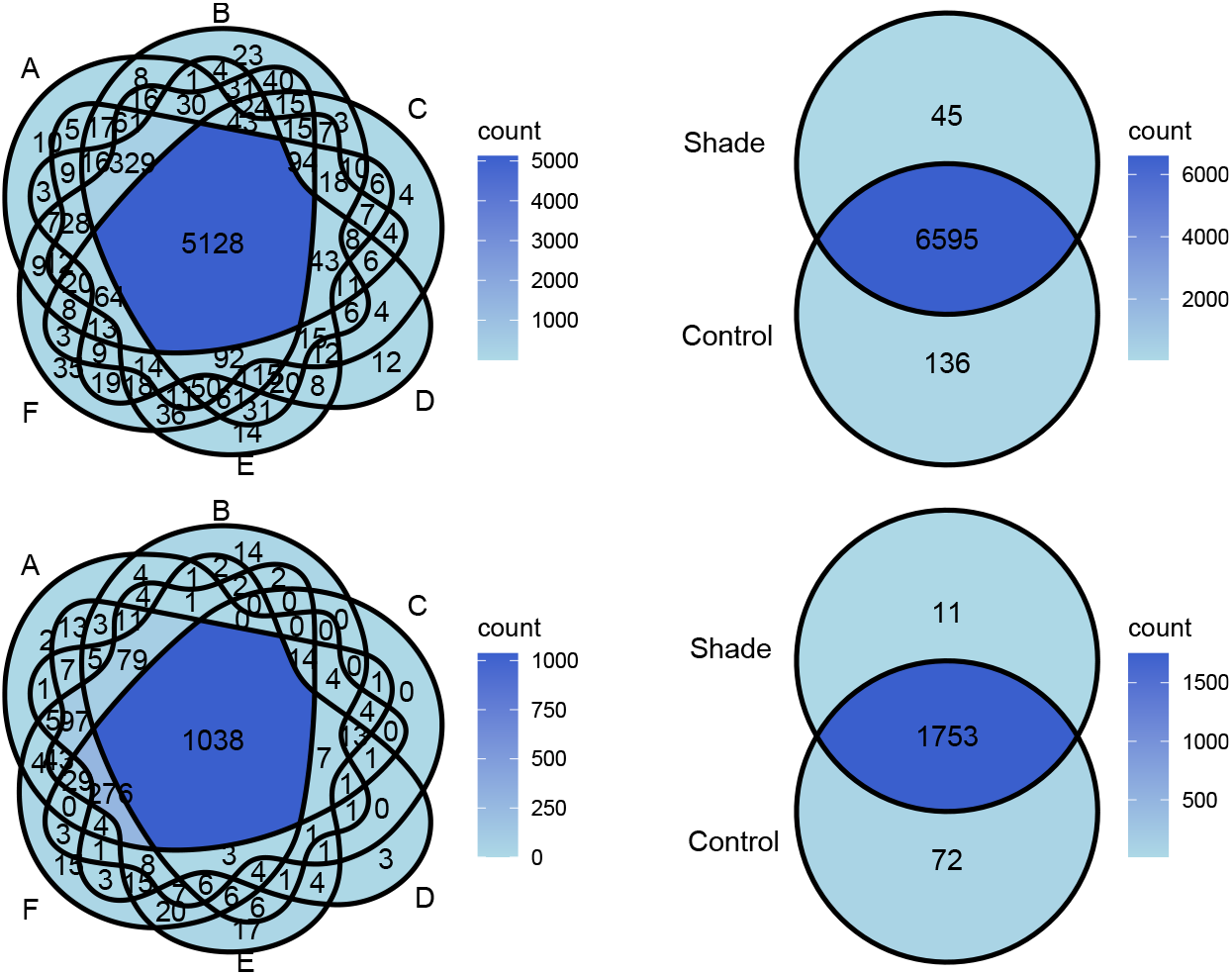
Venn diagrams showing unique and shared proteins among; top-left: *M. capitata* coral colonies, top-right: *M. capitata* colonies growing in control or shade treatment, bottom-left: zooxanthellae from each colony and bottom-right: zooxanthellae from colonies growing in control or shade treatment.

**Fig. S6.**
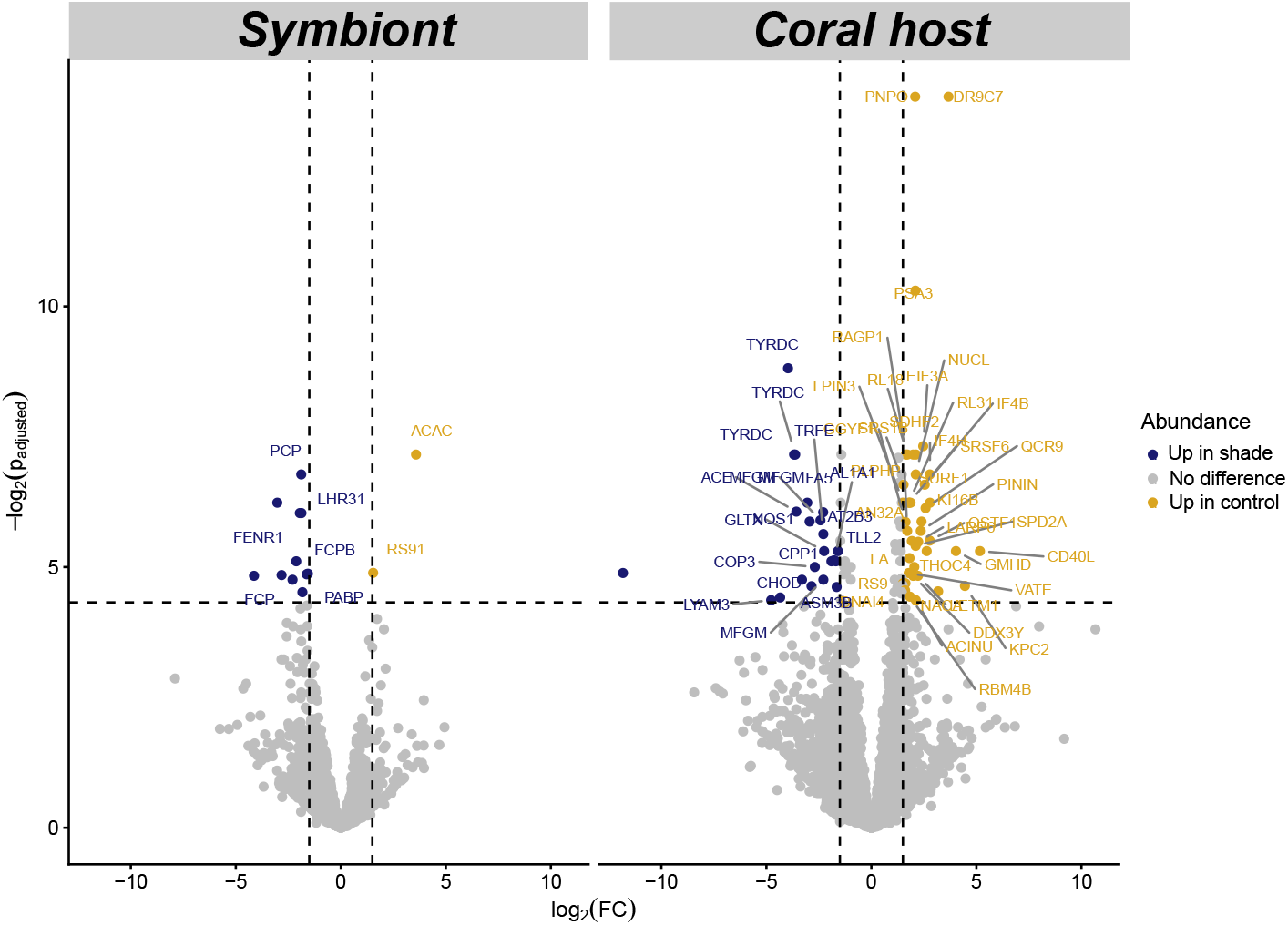
Volcano plots showing the results of differential protein abundance analysis in *Montipora capitata* and Symbiodiniaceae under control and and shaded conditions, along with the names of differentially abundant proteins retrieved from EggNOG-mapper. Dashed lines represent significance thresholds (*p*_adjusted_ *<* 0.05 and | log_2_ FC| *>* 1.5).

**Table S1.**
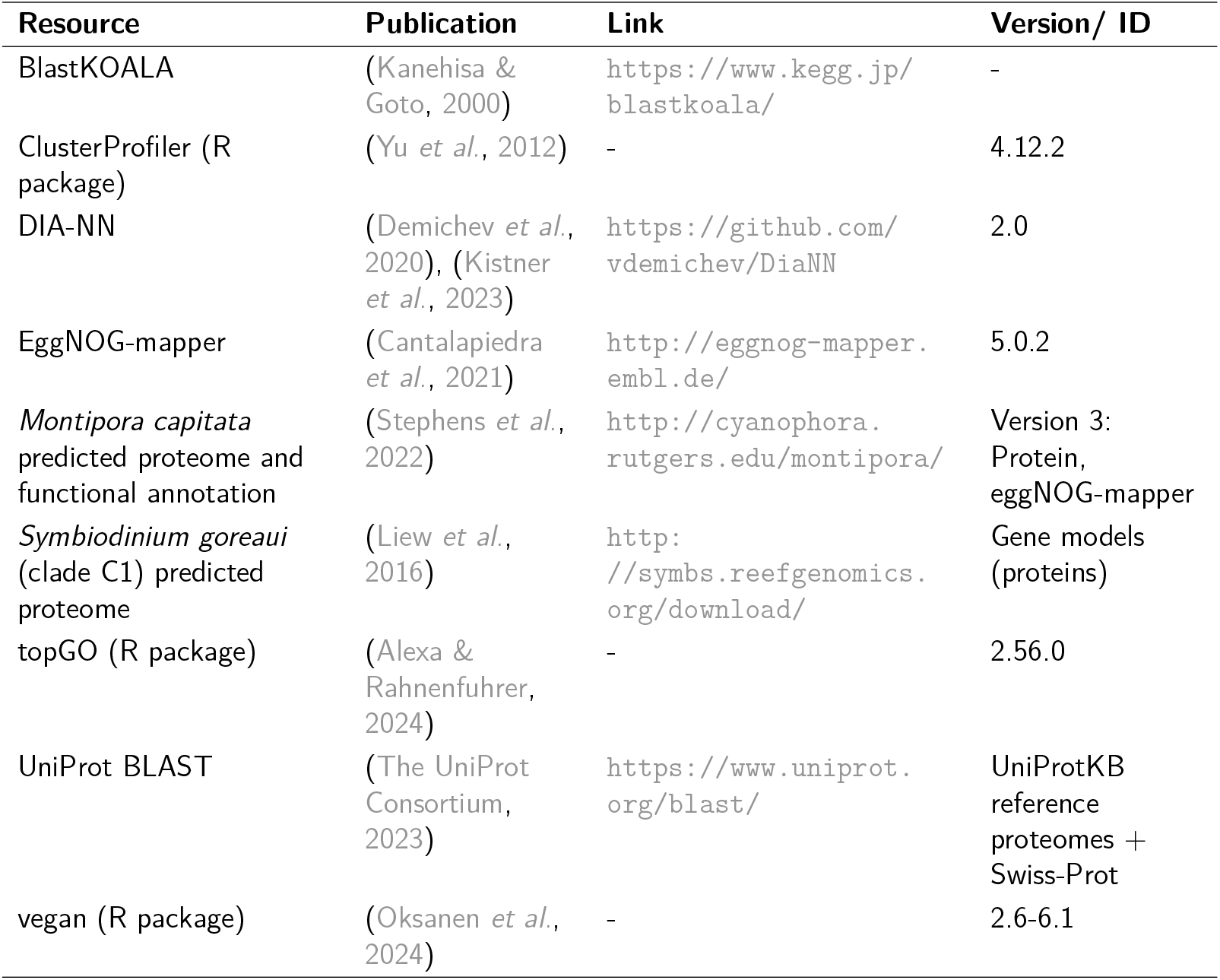
Key resources.

